# Dispersion of functional gradients across the lifespan

**DOI:** 10.1101/2020.02.27.968537

**Authors:** Richard A.I. Bethlehem, Casey Paquola, Jakob Seidlitz, Lisa Ronan, Boris Bernhardt, Cam-CAN Consortium, Kamen A. Tsvetanov

**Author notes:** Equal contribution (Richard A.I. Bethlehem, Casey Paquola).

## Abstract

Ageing is commonly associated with changes to segregation and integration of functional brain networks, but, in isolation, current network-based approaches struggle to elucidate changes across the many axes of functional organisation. However, the advent of gradient mapping techniques to neuroimaging provides a new means of studying functional organisation in a multi-dimensional connectivity space. Here, we studied ageing and behaviourally-relevant differences in a three-dimensional connectivity space using the Cambridge Centre for Ageing Neuroscience cohort (n=643). Building on gradient mapping techniques, we developed a set of measures to quantify the dispersion within and between functional communities. We detected a strong shift of the visual network across the lifespan from an extreme to a more central position in the 3D gradient space. In contrast, the dispersion distance of transmodal communities (dorsal attention, ventral attention, frontoparietal and default mode) did not change. However, these communities were increasingly dispersed with increasing age, reflecting more dissimilar functional connectivity profiles within each community. Increasing dispersion of frontoparietal and ventral attention networks, in particular, was associated negatively with cognition, measured by fluid intelligence. By using a technique that explicitly captures the ordering of functional systems in a multi-dimensional hierarchical framework, we identified behaviorally-relevant age-related differences of within and between network organisation. We propose that the study of functional gradients across the lifespan could provide insights that may facilitate the development of new strategies to maintain cognitive ability across the lifespan in health and disease.

## Introduction

With the increasing proportion of older adults in the worldwide population (Beard et al., 2016) and increasing burden of dementia in ageing societies, there is a pressing need to understand the neurobiology of cognitive ageing. Normal ageing is related to cognitive decline (Hedden and Gabrieli, 2004) and large-scale changes in brain dynamics (Li et al., 2015; Andrews-Hanna et al., 2007). Ageing is consistently related to weakened within-network connectivity co-occurring with increased between-network connectivity (Geerligs 2015, Betzel 2014, Chan 2014). Crucially, the coordination of within- and between-network connectivity supports the maintenance of general cognition (Tsvetanov et al., 2016) and performance in specific cognitive domains (Chan et al., 2014; Tsvetanov et al., 2018) across the lifespan.

Despite a large body of research on the decline and maintenance of brain network integrity during healthy ageing many inconsistencies remain. Most studies report both broad patterns of increased and decreased connectivity (Geerligs et al., 2015; Betzel et al., 2014; Chan et al., 2014), although others also report more one-directional in- (Ferreira et al., 2016) or decreases (Onoda et al., 2012; Damoiseaux et al., 2008). Thus, it is still unclear whether the observed age differences in functional network connectivity reflect a single global pattern or multiple patterns across cortical network hierarchical organization. In addition, the presence of multiple underlying connectivity patterns begs the question whether these patterns are uniquely or interactively contributing to maintaining cognitive function across the lifespan. Furthermore, standard network connectivity approaches, unlike data-driven and generative-modelling counterparts, are unable to disentangle between different physiological processes leading to the connectivity patterns (e.g. neuronal versus vascular origins of the fMRI BOLD signal) which are differentially affected by ageing (Tsvetanov et al., 2019). Given the importance of relative changes in functional connectivity to ageing and cognitive decline, a novel data-driven multi-dimensional framework approach is needed to directly address the behavioral relevance of age-related changes in large-scale organization. Here, we sought to directly address the behavioural relevance of age-related differences in large-scale organisation of resting-state functional connectivity with a novel multi-dimensional framework.

Recent studies of cortical dynamics and meta-analytic decoding of cognitive states have highlighted how brain regions participate in a range of functions (Braga and Leech, 2015; Yarkoni et al., 2011; Genon et al., 2018). Therefore, capturing the full extent of a region’s participation in the functional organisation of the cortex may require a more nuanced approach than comparison of connectivity within and between static functional communities. Embedding approaches have emerged as a tool to map a brain region’s functional relationships from resting state fMRI connectivity (Margulies et al., 2016; Tian et al., 2020; Haak et al., 2018). These data-driven techniques reveal multiple dimensions of cortical organisation, allowing each region to be characterised according to its position along many large-scale functional gradients. The alignment of the dominant functional gradients, sensory-to-transmodal and visual-to-somatomotor (Margulies et al., 2016), with the seminal model of the cortical hierarchy (Mesulam, 1998) supports their utility in understanding the fundamental principles of the cortical landscape in humans.

In order to assess age-related changes to the hierarchical functional organisation of the cortex, we first investigated how the principal gradients of resting-state connectivity change across the lifespan. These one-dimensional gradients aim to capture differentiation between systems which to some extent encapsulates both within as well as between network connectivity. For example, an increased bi-modality would indicate a segregation of the gradient extremes as well as an increased within-anchor connectivity pattern. We expected the principal gradient to recapitulate the sensory-to-transmodal organisation seen in previous studies (Margulies et al., 2016; Paquola et al., 2019), and hypothesised that age-related changes would be centralised in the default mode network, in line with previous studies of functional connectivity (Chan et al., 2014). Specifically, decreased segregation might be reflected on the principle gradient axes by a reduction of the range or bimodality of the gradient distribution showing that the anchors get closer. We also expected the second and third gradients would illustrate functional differentiation of visual, somatomotor and attention-related regions (Margulies et al., 2016; Paquola et al., 2019). Age-related changes within these functional systems have not previously been reported, though one meta-analysis of activation changes in healthy ageing identified hypo-activation in the visual cortex (Li et al., 2015).

To overcome the aforementioned challenges in neurocognitive research we further extended this existing gradient framework. Specifically, by building on gradient mapping techniques, we developed a set of measures to quantify the dispersion within and between functional communities in a multi-dimensional connectivity space. In particular, we sought to assess how functional communities shift in the multi-dimensional gradient space, whether these changes recapitulate findings using base functional connectivity, and finally whether these differences are of behavioural relevance in terms of cognitive decline. Predictions across multi-dimensional gradients are motivated by prior related work on network integration and segregation, which would suggest broadly reduced segregation between networks, particularly in the default mode and fronto-parietal networks (Chan et al., 2014; Geerligs et al., 2017; Betzel et al., 2014).

## Methods

### Dataset

The present study included 643 participants from the Cambridge Centre for Aging and Neuroscience (Cam-CAN) (Shafto et al., 2014). These participants were cognitively healthy adults (age range = 18-88) recruited from the local community. Ethical approval for the Cam-CAN study was obtained from the Cambridgeshire 2 (now East of England–Cambridge Central) Research Ethics Committee. Participants gave written informed consent. A subset of 514 individuals, previously described (Ronan et al., 2016), was used as the primary cohort where individuals’ Freesurfer reconstructions (see below) had already been processed and individually evaluated. The remaining data (n = 129, matched on age and sex) served as a hold-out dataset (Table S1) used for alignment of the diffusion map embeddings of the primary cohort to ensure the gradient maps were aligned across subjects.

**Table 1:** Demographics and in-scanner motion (average framewise displacement) of the primary cohort. SMD: standardised mean difference.

|  | F | M | SMD |
| --- | --- | --- | --- |
| n | 260 | 254 |  |
| age (mean (SD)) | 52.8 (18.68) | 53.32 (18.22) | 0.03 |
| motion (mean (SD)) | 45.51 (0.11) | 45.53 (0.11) | 0.16 |
| cattell (mean (SD)) | 31.68 (6.45) | 32.71 (6.78) | 0.16 |

### MRI acquisition and preprocessing

Structural and functional images were acquired on a 3T Siemens TIM Trio system employing a 32-channel head coil. A high resolution 3D T1-weighted structural image was acquired using a MPRAGE sequence (Repetition Time = 2250ms; Echo Time=2.99ms; TI=900ms; flip angle=9°; FOV=256mm × 240mm × 192mm; 1mm isotropic voxels; GRAPPA acceleration factor=2; acquisition time=4min 32s). A functional echo planar imaging scan was acquired while participants rested with eyes closed for 8min 40s (TR=1970ms; TE=30ms; flip angle=78 °; FOV = 192mm x 192mm; 32 axial slices; voxel size = 3mm x 3mm x 4.44mm). Data were quality-control checked by semiautomated scripts monitored by the Cam-CAN methods team (Taylor et al., 2017). Functional images underwent intra-modal and ICA-based motion correction using ICA-AROMA (Pruim et al., 2015). This included, removal of the first 5 volumes, framewise displacement and spike regression, spatial smoothing using a 2mm gaussian kernel, co-registration between T1-weighted and functional images and finally removal of motion artefacts using automated ICA-AROMA. Denoised BOLD timeseries were projected from volumetric native space to subject-specific cortical surfaces, which were rendered from the T1-weighted scans using Freesurfer v5.2 (Fischl, 1999; Dale, 1999).

Second level analyses were conducted at surface level using MATLAB and SurfStat (Worsley et al., 2009). We averaged pre-processed vertex-based time-series within 1012 equally sized, spatially contiguous nodes (Hong et al., 2017), then used pairwise Pearson correlation and z-standarisation to create individual functional connectivity matrices. In line with previous studies assessing gradients in young adults (Paquola et al., 2019; Vos de Wael et al., 2018; Margulies et al., 2016) connectivity matrices were subjected to row-wise thresholding (top 10% of edges maintained), then converted into a normalised angle matrix. Diffusion map embedding, a non-linear dimensionality reduction technique (Coifman and Lafon, 2006), was employed to resolve the gradients of subject-level connectomes. A group-average functional connectivity matrix constructed using the hold-out sample underwent the same diffusion map embedding procedure and were reordered to conform with the conventions set out in Margulies et al., (2016). The first three gradients, explaining 52% of variance in the affinity matrix of the hold-out cohort (Supplementary Material: Table S1 and Figure S1A and S3), showed functional differentiation running from sensory-to-transmodal (G1), visual-to-insula (G2) and somatomotor-to-insula (G3)(Figure S1A). Together, these gradients describe functional discrimination between sensory modalities and across levels of the cortical hierarchy, (ie: sensory processing, attentional modulation then higher-order cognition). Individual embedding solutions were aligned to the group-level hold-out embedding via Procrustes rotations (Langs et al., 2015; Wang and Mahadevan, 2008). The Procrustes alignment enables comparison across individual embedding solutions, provided the original data is equivalent enough to produce comparable Euclidean spaces (Coifman and Hirn, 2014).

### Age-related differences in gradient values

In our primary analyses, we assessed age-related differences in nodal gradient values by fitting linear models, controlling for head motion and sex. To synoptically visualise these age related effects we discretised the hold-out gradient into 10 equally sized bins of the gradient values and calculated the average and standard deviation of t-statistics from the linear model in each bin. Post-hoc exploratory analyses focused on summary measures derived from the gradient distributions and included the range of the gradient distribution and its bimodality as these were hypothesized to be indicative of changes to the gradient anchors. The latter was quantified using Hartigan’s dip test (Hartigan and Hartigan., 1985) (see supplemental Figure 1 and 2, and supplemental table 2).

### Age-related differences in multi-dimensional gradient dispersion

To investigate multi-dimensional differences in cortical organisation, we quantified a new metric, termed here “*dispersion*” of established functional networks (Yeo et al., 2011) in 3D space. Each axis of this 3D space was defined by the values along the first three gradients. *Within network dispersion* was quantified as sum squared Euclidean distance of network nodes to the network centroid at individual level. *Between network dispersion* was calculated as the Euclidean distance between network centroids. These metrics were calculated for each subject within the individualised, aligned gradient space. We used linear models, correcting for head motion and sex, to estimate the association of age with network centroid position, within network dispersion and between network dispersion. For each model, we computed a null distribution of age-related differences using 1000 spherical rotations of the original parcellation scheme, thereby ensuring that for each permutation of the null distribution the spatial autocorrelation was retained (Alexander-Bloch et al., 2018).

### Alternative measures of brain function and structure

To determine the unique contribution of network dispersion versus previously reported estimates of unidimensional within network connectivity (WNC) (Yan et al., 2013), we ran robust linear regression (Huber and Ronchetti, 2009) for each Yeo network on dispersion controlling for within network connectivity of the same network (and vice versa) using the MASS library in R. For example, to assess the unique age-related contribution of network dispersion within the DMN we ran the following linear model (and vice versa for within network connectivity):

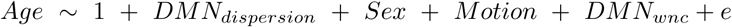

In addition, to determine whether the proposed multi-dimensional gradient approach captures similar aspects of network segregation and integration as edge-based methods, we calculated segregation [difference in within-vs between-network connectivity, proportional to within-network connectivity (Chan et al., 2014)] and normalised clustering coefficient (Rubinov and Sporns, 2010) for each node, then repeated the robust linear models estimating the effect of age. Robust linear regression was used to down-weight datapoints with large residuals which strengthens the idea that results are not driven by a small number of outlier datapoints or subjects (Huber and Ronchetti, 2009).

### Behavioural relevance of network dispersion

Finally, to determine whether age-related cognitive decline could be explained by the contribution of network dispersion, we conducted mediation analysis on the networks showing age-related differences on dispersion. For the purpose, cognitive function was assessed using the Cattell fluid intelligence scale (Cattell, 1963). This analyses was conducted in R using the mediation package (Tingley et al., 2014). Briefly, this approach implements bootstrapping to compute a point estimate of the indirect effect over 1000 bootstrap iterations so it is less sensitive to non-normally distributed mediators and smaller sample sizes (Hayes and Preacher, 2010; Preacher and Hayes, 2004).

To further assess the nature of the associations between cognition and gradient dispersion, i.e. whether it is age-dependent and/or age independent, we conducted a moderation analysis. Specifically, we constructed a multiple linear regression model where age, gradient dispersion and their interaction (age x dispersion) were used as independent variables and Cattell total scores were used as the dependent variable. Covariates of no interest in the model included sex and head motion. We visualised any potential moderating effects of network dispersion by plotting the relationship between Dispersion and Cattell score for three different age groups.

### Code and data availability

Data and code used in this manuscript are available from https://github.com/rb643/GradientDispersion.

## Results

### Multi-dimensional age-related differences

We aimed to quantify age-related differences in the global orchestration of functional connectivity by projecting established functional communities into the 3D gradient space. Functional communities were compactly localized in the group-average 3D gradient space (Figure 1A). The visual network occupied the most extreme, segregated position, driven by the second gradient. The default mode network also occupied an extreme position, driven by the first gradient, and was almost completely bordered from other networks by nodes from the frontoparietal network. Somatomotor and ventral attention networks populated one side of the 3D space, covering the range of the third gradient. Finally, dorsal attention and limbic networks resided close to the centre of space.

**Figure 1:**
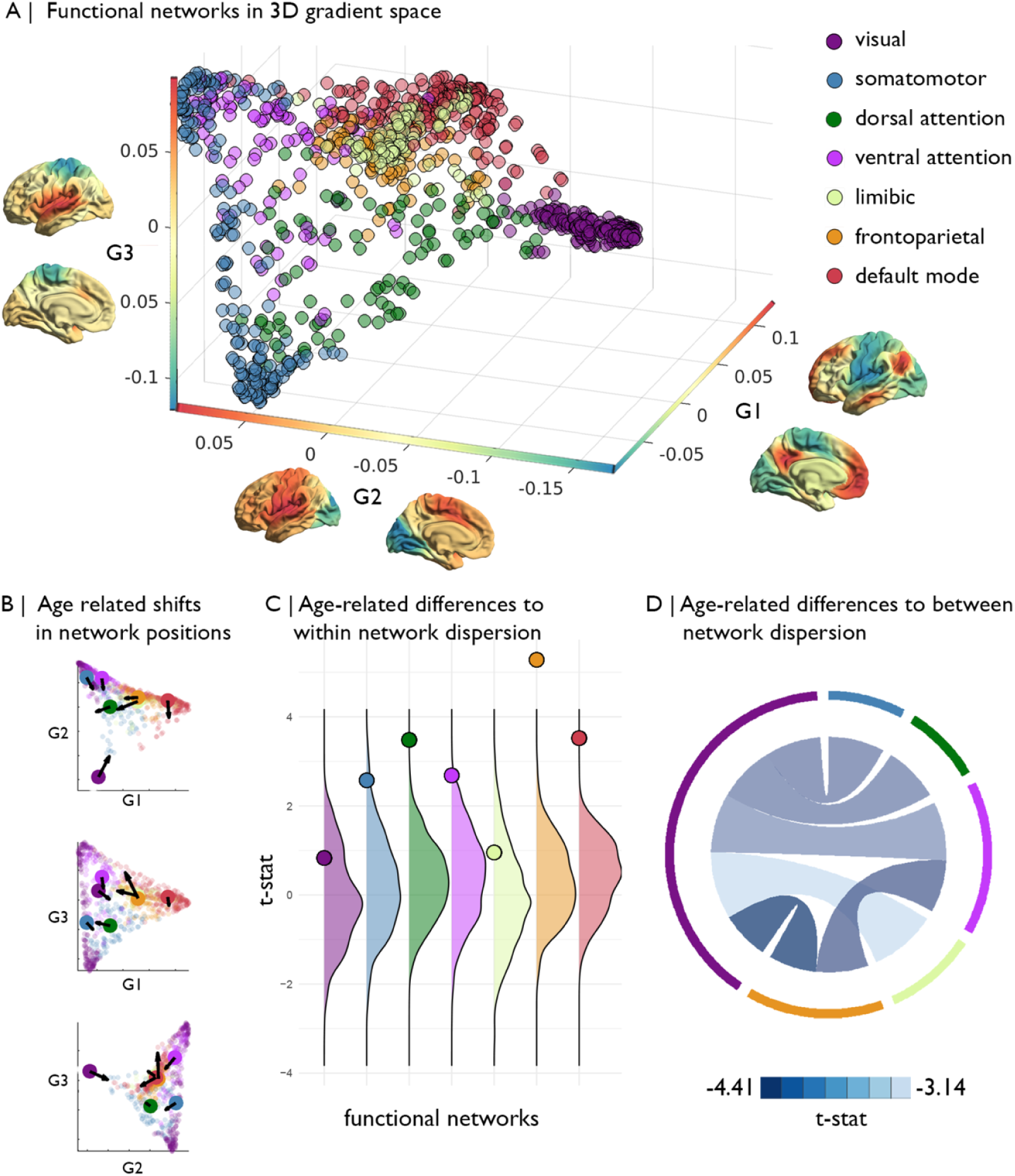
Functional gradients across healthy adulthood. A) First three gradients projected into a 3-dimensional gradient space and coloured by its Yeo network classification. Average gradient topology of the holdout are depicted on each axis. B) Age-related shift in the centroid of each Yep network in gradient space. Arrows reflect the direction of the centroid shift and are scaled by effect size. C) Age-related difference in within network dispersion. Dots indicate the actual t-statistic of the dispersion. Distribution capture the spin-permuted null distribution t-values. D) Age-related difference in between network dispersion. Only pairs showing a significant age-related change are depicted.

The visual network became less differentiated from other networks with increasing age, as evidenced by significantly decreasing *between network dispersion* (i.e. the distance between the centroids of each network) (4.35 *<*t*<*-2.88, p_spin_*<*0.068, Figure 1D). This age-related difference occurred relatively uniformly for all nodes of the visual network, as its *within network dispersion* was not related to age (t=-2.49, p_spin_=0.77). In contrast, the dorsal attention, ventral attention, fronto-parietal and default mode networks each became more dispersed in the 3D gradient space with increasing age (t=3.49/2.69/5.53/3.52, p_spin_ *<*0.05), reflecting more dissimilar functional connectivity profiles within the respective networks (Figure 1B & C). To explore whether age related differences in dispersion were driven by a particular gradient, we examined the correlation between dispersion and subjects individual principal component loadings for the principal component of each gradient (Supplementary material, Figure S5). There was no indication that dispersion in any of the Yeo networks was specifically or preferentially associated with one of the gradients (Supplementary Figure 5).

Notably, the overarching topology of the 3D gradient space was relatively stable across age. The strongest age-related differences were observed in the visual-to-insula gradient (G2) (Supplementary Figure 1 & 2). Gradient values of occipital areas significantly increased with age. In parallel, values in insula nodes, occupying the top of the gradient, decreased with age. Together the node-wise differences produced an overall reduction in the range of the gradient (t=-3.29, p_FDR_=0.0033) and more pronounced bimodality (t=5.18, p_FDR_*<*0.0001), representing reduced functional differentiation across the entire axis coupled with consolidation within each functional system. The apex of the principle gradient (G1) remained constant across the age range, however, prefrontal regions tended to shift down the gradient with increasing age. Prefrontal regions also shifted up the somatomotor-to-insula gradient (G3), showing a shift in functional affiliation from the default mode apex of G1 towards the attention focused apex of G3. Range and bimodality of G2 and G3 did not significantly change with age (Supplementary Table 2).

### Relation to alternative measures of brain function and structure

We hypothesised that the gradient dispersion technique offer a higher-dimensional account of functional reorganisation throughout the lifespan, and would thus partially overlap with findings using prebiously established measures of functional topology. Average within network functional connectivity was not associated with age, suggesting within network dispersion is more sensitive to lifespan changes (Figure 2A). Furthermore, age-related difference in within network dispersion were independent of within network connectivity (Figure 2B; Table 2). Within network dispersion showed more similar age-related differences to the clustering coefficient (Rubinov and Sporns, 2010) (Figure 2C), but these were not concordant with age-related difference to “segregation” (Chan et al., 2014; Zonneveld et al., 2019). The segregation metric highlighted increases in the limbic network at later ages (Figure 2C). In contrast, dispersion and clustering indicated the strongest differences in ventral attention, dorsal attention, fronto-parietal and default mode networks. Dispersion, specifically, demonstrated increased sensitivity for detecting age-related difference in fronto-parietal and default mode networks. In addition, between network dispersion, unlike between network connectivity, was sensitive to age-related differences (Figure 2D).

**Figure 2:**
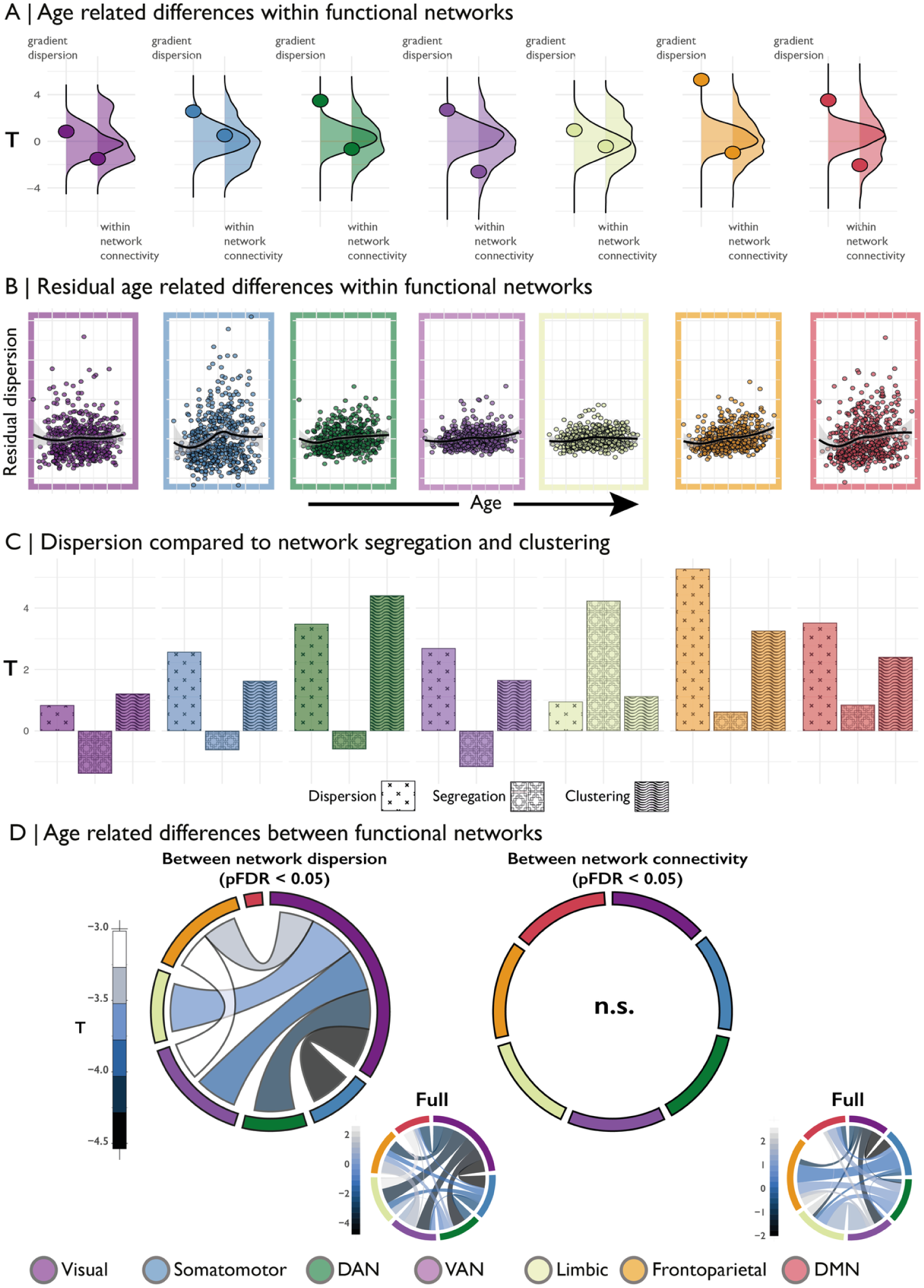
Panel A shows the age-related difference (t-statistic) in within network dispersion and within network connectivity for each of Yeo network. Underlying density plots show the null distributions of t-statistics derived from spin permutations. Panel B shows the residuals of the dispersion model (including controls for sex, motion and within network connectivity) against age residuals for the same model for each Yeo network. Panel C depicts t-statistics for age-related differences in Yeo network for within network dispersion, segregation (Chan et al., 2014) and clustering (Rubinov and Sporns, 2010). Panel D shows the between network dispersion and between network connectivity. Network borders are scaled according to the size of the total effect from that community (e.g. the visual network is largest in the left panel as most significant between network dispersion involved the visual network. Insert panels on the right show the full between network pattern for all connections including non-significant ones.

**Table 2:** Age related effects on dispersion and within network connectivity per Yeo community using robust linear regression controlling for motion, sex and opposite network metric.

| model | Withing Network Dispersion |  |  | Within Network Connectivity |  |  |
| --- | --- | --- | --- | --- | --- | --- |
|  | F | t | pFDR | F2 | t2 | pFDR |
| DMN | 28.89 | 5.38 | 4.09e-07 | 1.0 | -1.0 | 0.63 |
| Fronto Parietal | 50.21 | 7.09 | 3.24e-11 | 0.58 | 0.76 | 0.63 |
| Sensory Motor | 14.74 | 3.83 | 0.0 | 0.37 | 0.61 | 0.63 |
| DAN | 21.88 | 4.69 | 8.7e-06 | 0.1 | 0.32 | 0.75 |
| VAN | 15.73 | 3.96 | 0.0 | 4.64 | -2.17 | 0.22 |
| Limbic | 9.98 | 3.15 | 0.0 | 0.54 | -0.74 | 0.63 |
| Visual | 1.04 | 1.02 | 0.31 | 1.37 | -1.17 | 0.63 |

### Influence of cortical morphology

Ageing is strongly associated with cortical atrophy (Lowe et al., 2019). We performed surface based linear regression to assess the age-related differences in cortical thickness and detected widespread cortical thinning (Figure 3). To determine whether this thinning influence age-related differences to the gradient dispersion, we repeated unimodal gradient analysis and within network dispersion models while controlling for cortical thickness. Results revealed identical patterns of gradient changes after controlling for cortical thickness. We found that controlling for cortical thinning slightly increases the linear effect of age on network dispersion as all previously significant networks remain significant and show a slight increase in F and T-values.

**Figure 3:**
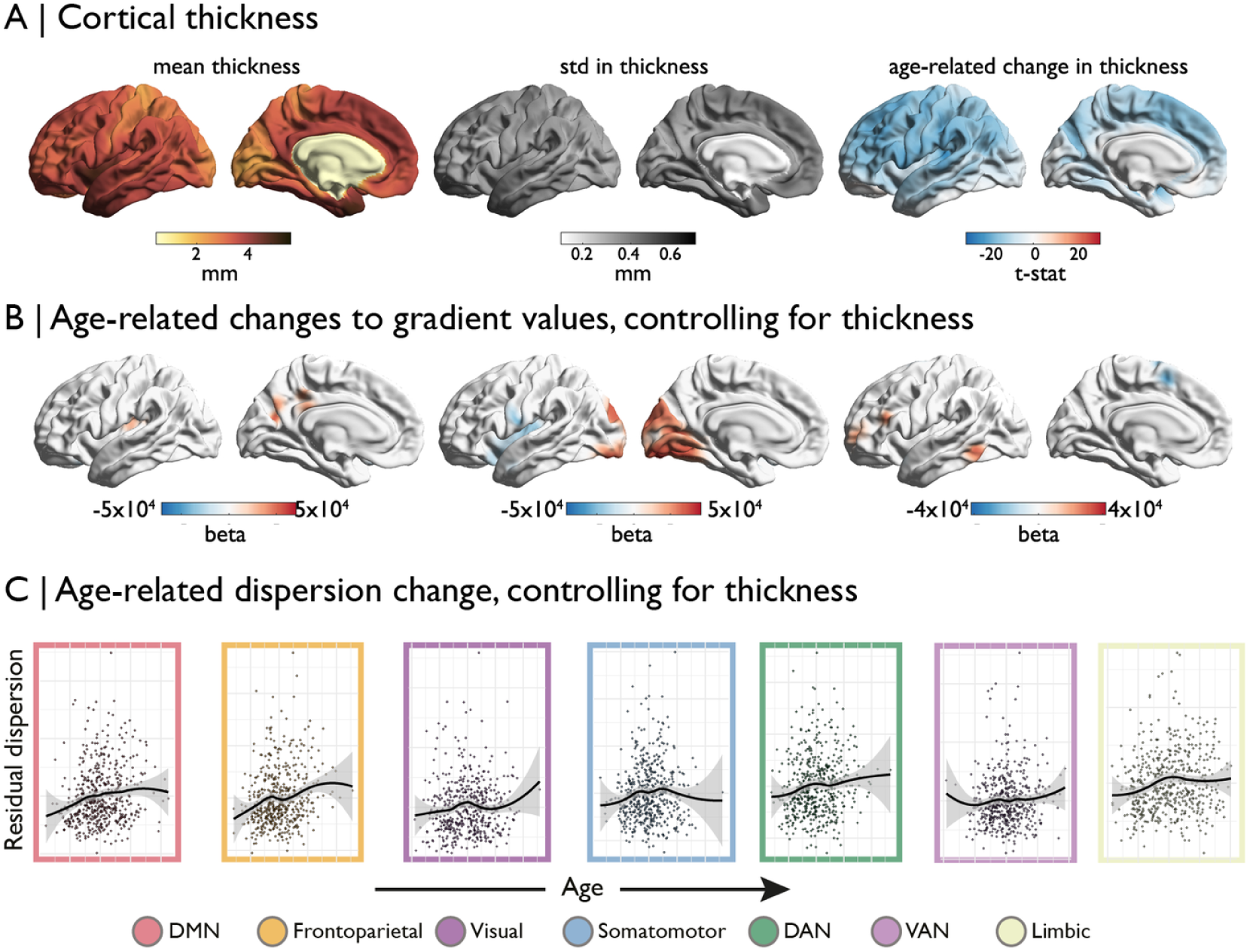
Morphological changes in cortical thickness are widespread and uniformly negative (Panel A). Age-related changes to the gradients are not effected by this cortical atrophy (Panel B) nor are the multidimensional changes in gradient dispersion (Panel C).

### Relation to cognition

Finally, we explored whether the observed age-related differences in network dispersion were behaviorally relevant, as assessed with the Cattell fluid intelligence scale (Cattell, 1963). Of the four networks that showed an age-related change in dispersion using spin permutations (dorsal attention, ventral attention, fronto-parietal, default mode) only the ventral attention network and the fronto-parietal network showed a small but significant average causal mediation effect (ACME) on age-related decreases in fluid intelligence (Figure 4A-D). The dorsal attention network also showed a small significant ACME (p = 0.032). Both the fronto-parietal and ventral attention network show a similar directionality when visualised with a median split. Specifically, both show that individuals with lower dispersion are likely to maintain higher cognitive scores later in life. Unimodal gradient changes (Figure S1) did not appear to moderate or mediate the relationship between age and cognition, however (Supplementary Materials, Figure S4).

**Figure 4:**
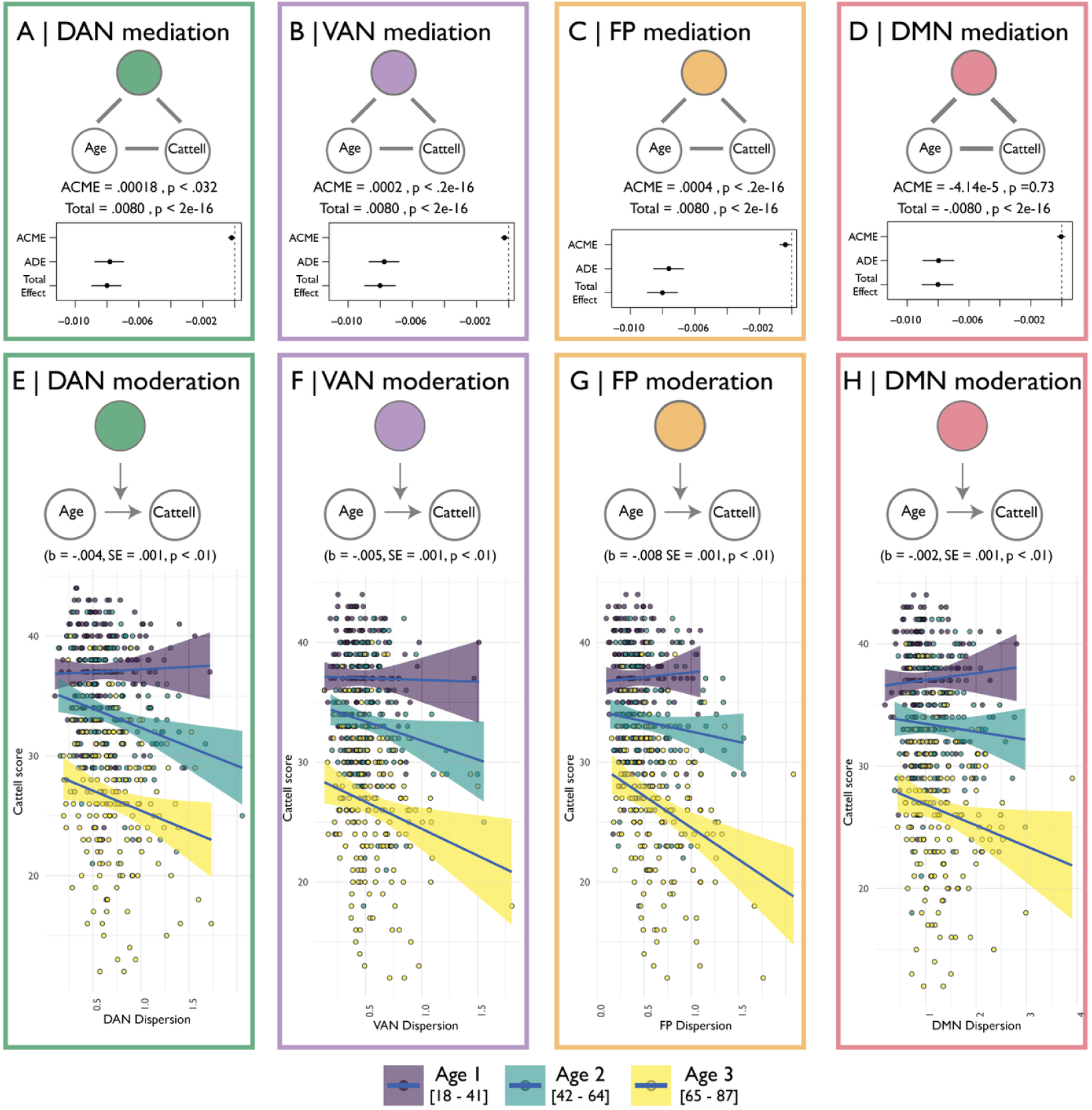
Mediation of age-related cognitive decline. Panels A-D show the mediation effects of network showing significant age-related change in gradient dispersion on the age-related change in cognitive decline. For each network the path and average causal mediation effect (ACME) based on bootstrapping are shown. Dot-plots also provide the average direct effect (ADE) estimates and confidence intervals. Panels E-H show the moderating effects of the same networks’ dispersion on age and Cattel score. Scatterplots show how this effect varied in three bins of the total age-range.

## Discussion

We demonstrate that the ordering of regions within large-scale functional gradients remains relatively stable across the lifespan, but the dispersion of fronto-parietal and ventral attention networks in a 3D gradient space was related to cognitive decline. Decreasing dispersion was not preferentially associated with one gradient dimension, indicating the importance of multiple global patterns of connectivity for maintaining performance in old age. Our findings bolster existing literature on the importance of network integration and segregation for cognitive function (Geerligs et al., 2017; Andrews-Hanna et al., 2007; Betzel et al., 2014; Geerligs et al., 2014; Chan et al., 2014; Li et al., 2015; Tsvetanov et al., 2016; Zonneveld et al., 2019). By using a technique that explicitly captures the ordering of functional systems in a hierarchical framework, we exposed subtle age-related differences in within and between network organisation that conventional linear interactions did not detect and that was robust against known changes in cortical morphology. Furthermore, these effects were behaviourally relevant as they explained partly the relationship between age and fluid intelligence.

At a uni-dimensional level, we also observed age-related increases in the gradient values of the visual network. This indicated that, on the visual-insular axes of the gradient topology, the visual cortex became more similar to the rest of the cortex. This was captured by an overall decreased range in the gradient values and an increase of gradient values particularly in visual cortex. This de-differentiation of the visual cortex was also apparent in the context of the multi-dimensional gradient space where we observed that nodes within the Yeo visual network cluster almost uniformly moved closer towards the centre of all the other networks. Prior literature has indicated that in older adults the visual network shows signs of hyperactivity combined with decreased activation in control and default mode networks (Li et al., 2015). Interestingly, while we observed a de-differentiation of the visual cortex, we also observe an increased dispersion of frontal and default mode network that might fit with this prior finding of decreased activity in control networks. Thus, at a unimodal gradient level, we observed increased indicators of segregation within the visuo-insular gradient (e.g. increased bimodality).

Within specific networks we observed the strongest increased dispersion in default mode and fronto-parietal networks, but also in the ventral and dorsal attention networks as well as the somato-motor network. Increased dispersion is captures decreased similarity across multiple gradient domains. This notion is well aligned with prior reports of broad decreased functional connectivity (Zonneveld et al., 2019) and broad decreased functional cohesion (Betzel et al., 2014). Comparing dispersion in multi-dimensional gradient space to other approaches of measuring network changes such as clustering (Betzel et al., 2014), within network connectivity (Zonneveld et al., 2019) and segregation (Chan et al., 2014) more directly, we observe similar patterns of decreased between network connectivity, yet gradient dispersion seems to have an overall increased sensitivity to age-related differences. These age-related differences in dispersion also proved to be significant mediators and moderators of the negative association between age and fluid intelligence as measured by the Cattell (Cattell, 1963). Qualitative inspection of this effect suggested that this increasingly plays a role with increased age. In other words, maintaining network topology in older age may be a key component to maintaining healthy cognition.

In addition, formal testing of the moderation effects suggested that maintaining youth-like network dispersion increasingly plays a role with increased age. Our findings are consistent with previous reports based on neuronal signatures from magnetoencephalography data (Bruffaerts et al., 2019) or haemodynamic signatures from fMRI BOLD data only after accurate modelling of physiological and vascular differences (Geerligs et al., 2017; Tsvetanov et al., 2016). This suggests that gradient-based estimates might reflect more closely signals of neuronal origin, compared to standard estimates of hemodynamic connectivity, and confirm that maintaining network topology in older age may be a key component to maintaining healthy cognition.

Some caveats to the present study should be noted however. First, we currently used a single cognitive measure to determine behavioural relevance (e.g. Cattell). Future studies will need to investigate the behavioural relevance of gradient dispersion across other cognitive domains. Second, the present results are based on a population-based cross-sectional cohort, and cannot directly speak to individual subjects’ changes over time. The above discussion of age effects is therefore restricted to the effects of age and its correlates, as assessed across individuals, rather than the dynamic process of individual ageing ageing per se. Future longitudinal studies are required to confirm whether the gradient dispersion is sensitive to detect changes in functional reorganisation reorganisation in individual progression either in healthy or diseased state. Finally, in the current study a hold-out dataset was used for procrustes alignment of the diffusion map embeddings to ensure accurate gradient map alignment across individuals. Future studies could consider the use of existing cohort-matched gradient map alignments. For that purpose the repository containing the code used in the present analysis also includes the individual matrices and code used to generate the hold-out alignment sample. Future studies could use this to generate their own out of sample reference alignment to match specific demographic factors (e.g: age and/or sex).

In sum, we show studying functional organisation in a multidimensional gradient framework can provide increased sensitivity to capturing and furthering previously established age-related changes in functional network topology. They are robust to potential confounds of known morphological changes and the captured change in topology has a significant mediating effect on age-related cognitive decline. The approach and the findings have implications for preventative and interventional strategies that aim to target maintenance of multidimensional network organisation to promote mental wellbeing in health and disease

## Acknowledgements

This project would not have been possible without the work already done by the Cam-CAN consortium and their open sharing of data with the scientific community.

