## Supplemental information for "Dispersion of functional gradients across the lifespan"

[1]University of Cambridge, Department of Psychiatry, Brain Mapping Unit
[2]* author contributed equally
[3]McConnell Brain Imaging Centre, Montreal Neurological Institute and Hospital, McGill University, Montreal, QC, Canada;
[4]National Institutes of Health
[5]University of Cambridge

February 19, 2020

### Holdout sample characteristics

| | FEMALE | MALE | p | test |
|---|---|---|---|---|
| n | 64 | 65 | | |
| Age (mean (SD)) | 57.88 (19.23) | 61.49 (17.83) | 0.27 | |
| Cattell_TotalScore (mean (SD)) | 30.78 (7.33) | 30.13 (7.06) | 0.62 | |

Table 1: Demographics for the holdout sample used for realignment of the gradient.

### Age-related differences in gradient values

In our primary analyses, we assessed age-related differences in nodal gradient values by fitting linear models, controlling for head motion and sex. To synoptically visualise these age related effects we discretised the hold-out gradient into 10 equally sized bins of the gradient values and calculated the average and standard deviation of t-statistics from the linear model in each bin. Post-hoc exploratory analyses focused on summary measures derived from the gradient distributions and included the range of the gradient distribution and its bimodality as these were hypothesized to be indicative of changes to the gradient anchors. The latter was quantified using Hartigan's dip test (Hartigan and Hartigan, 1985) (see supplemental Figure 1 and supplemental table 2).

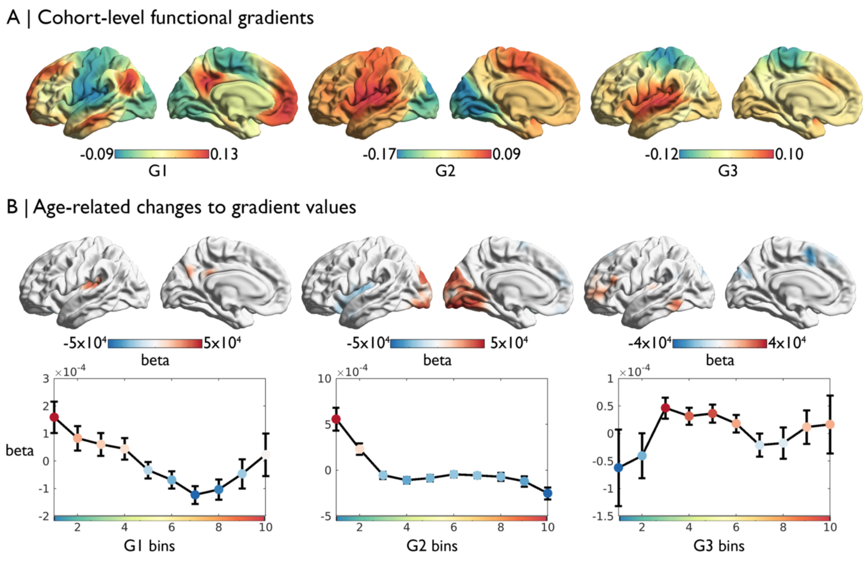


Figure 1: Unidimensional age-related changes. A) The first three principal gradients depict transitions from sensory-transmodal (G1), visual-insula (G2), and somatomotor-insula (G3). B) Beta estimates of age-related differences in gradient values, shown on the cortical surface with p<0.0083 threshold, and averaged within ten discrete bins of the gradient with standard error bars. C

### Range and bimodality

To assess univariate changes of the individual gradient cross-sectionally we quantified the range and bimodality of the gradient distributions. Bimodality was quantified using Hartigans dip-test of unimodality (Hartigan and Hartigan, 1985). Age-related changes were assess using linear models correcting for sex and head motion.

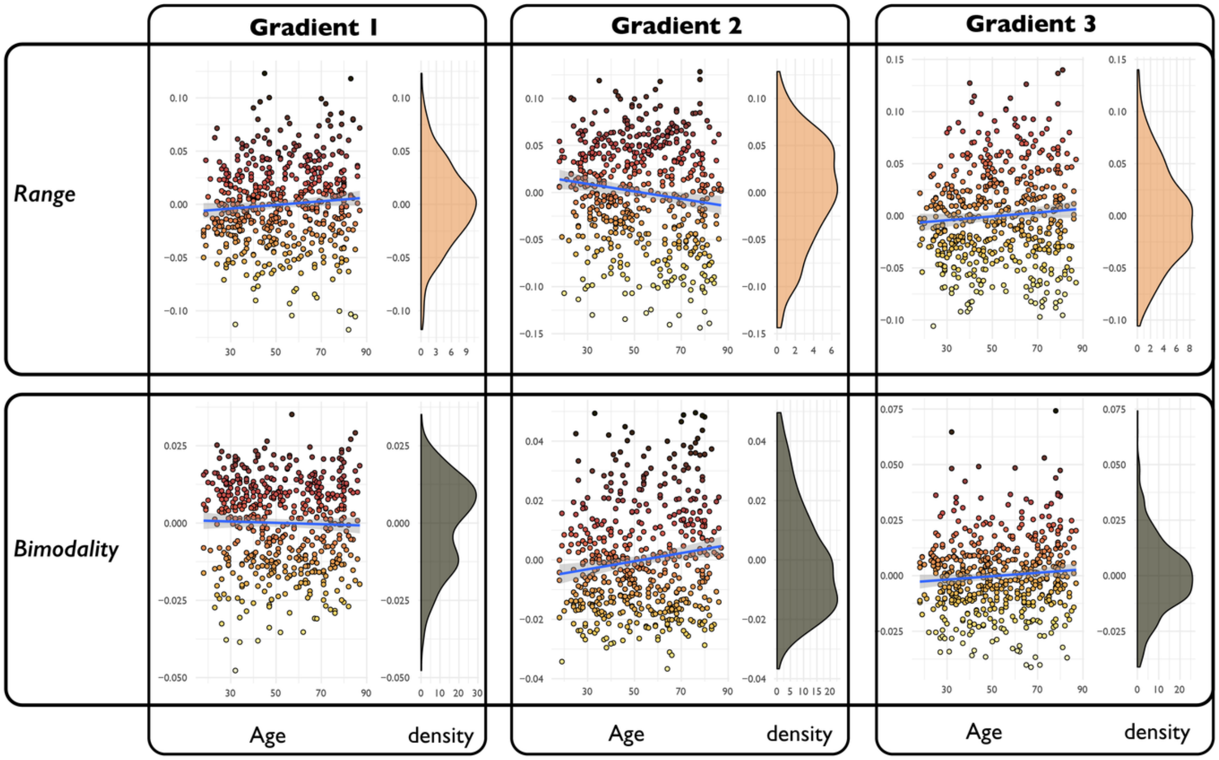


Figure 2: Residual range and bimodality of gradient distributions corrected for motion and sex plotted against age. Age-related changes from the same linear model are shown in Table 2.

| | | beta | se | t | pfdr |
|---|---|---|---|---|---|
| range | G1 | 0.0 | 0.0 | 2.01 | 0.09 |
| | G2 | -0.0 | 0.0 | -3.29 | 0.0 |
| | G3 | 0.0 | 0.0 | 1.85 | 0.1 |
| hartigan's dip | G1 | 0.01 | 0.01 | 1.01 | 0.38 |
| | G2 | 0.05 | 0.01 | 5.18 | 0 |
| | G3 | 0.01 | 0.02 | 0.57 | 0.57 |

Table 2: T-values from age-related changes on range and bimodality corrected for sex and head motion.

### Alignment and variance explained

To determine successful alignment between the main and holdout datasets we explored the amount of variance explained by individual eigenvectors pre- and post Procrustes alignment (Langs et al., 2015; Wang and Mahadevan, 2008). Procrustes alignment leads to a substantial shift in the matching. In the main manuscript the first and second eigenvectors are described as the second and first gradient, respectively, to align with seminal literature (Margulies et al., 2016) and because the sensory-transmodal gradient was detected in all subjects, unlike the visual-to-insula gradient.

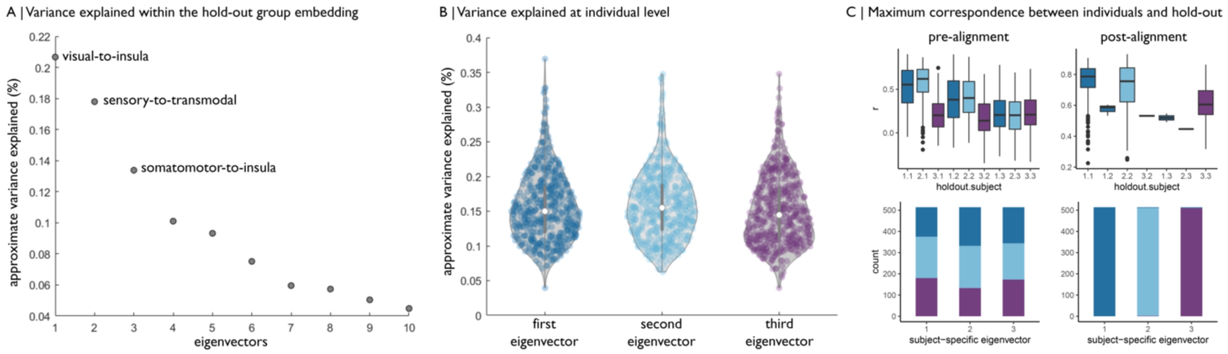


Figure 3: Variance explained in normalised affinity matrices by the first three eigenvectors. (A) Scree plot depicting the approximate variance explained within the hold-out group. (B) Violin plots show the variance explained by eigenvectors in individual subjects prior to Procrustes alignment to the hold out embedding. (C) We identified the most similar gradients in holdout and single subjects using Pearson correlations between the top three eigenvectors. The boxplots show the spatial correlation between the hold out and single - subject eigenvectors, with the factors representing which hold out eigenvector was maximally similar to the single-subject eigenvector. Ie: 1.2 represents the match of the first hold out eigenvector with the subject's second eigenvector. Below, the stacked columns show the proportion of each subject's eigenvectors that were matched to each hold out eigenvector.

### Mediation and moderation of unimodal gradients on age and cognition

To test the specificity of our dispersion measures we also conducted the same mediation and moderation analyses on the mean gradient values for the significant cluster on gradient. Gradient values on the second gradient did not reveal a moderating nor a mediating effect on the relationship between age and Cattell (Figure 4).

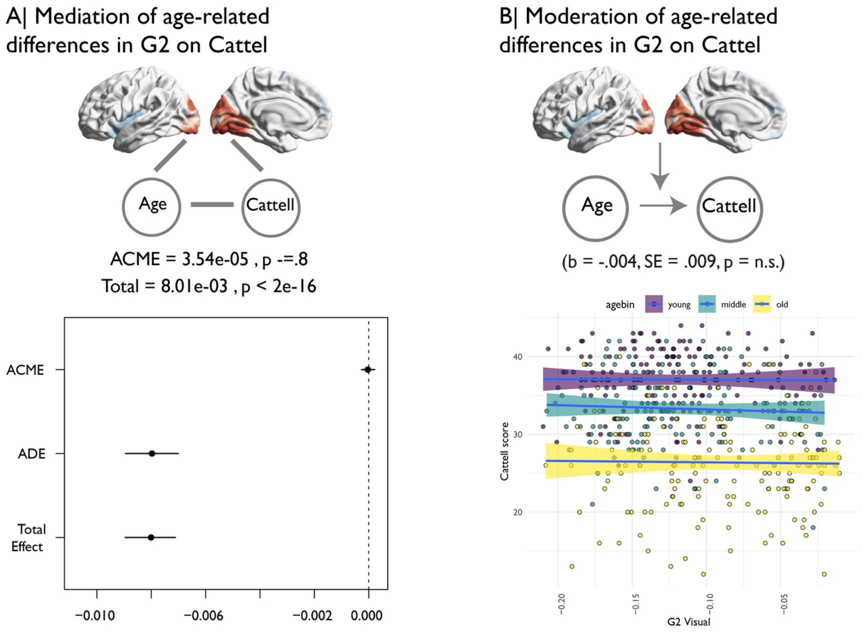


Figure 4: Mediation and moderation analysis of gradient values on age and Cattell

### Relation between dispersion and individual gradients

In order to delineate whether individual gradient contributed disproportionally to the Yeo network specific dispersion we quantified the contribution by looking at the correlation between the first principal component of each gradient and the the dispersion of each Yeo community. Although the gradient principal component do not correlate across all network there also was no clear evidence of a singular preferential association for a particular network.

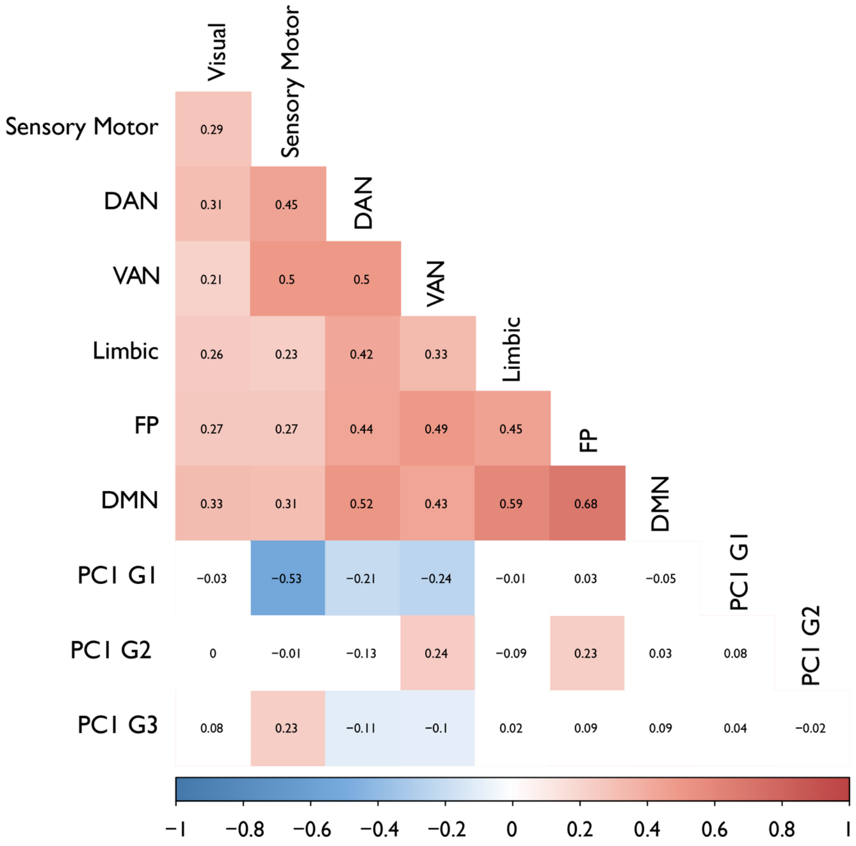


Figure 5: Dispersion to gradient PC correlations. Correlations with an uncorrected $p < 0.05$ are coloured according to their sign.
